# Strategies to improve neurite outgrowth from primary neurons in gelatin methacrylate hydrogels polymerized with visible light exposure inside a microscale 3D model

**DOI:** 10.64898/2026.09.28.755099

**Authors:** Iryna Liubchak, Joseph Sadden, Tanya Solomon, Yvonne Xie, Sai Cheong Ng, Tanya Bennet, Nasrin Zohreh, Peter Van den Doel, Alex Pieters, Paul Juralowicz, Alexis Hilts, Alicia Fung, Samantha Mung, Kaiwen Liu, Tara M. Caffrey, Karen C. Cheung

## Abstract

Gelatin methacrylate (GelMA) hydrogels offer many advantageous properties such as excellent biocompatibility, tunable stiffness, and rapid fabrication, making them attractive materials for neural systems-on-a-chip devices used to study axonal outgrowth in three-dimensional (3D) environments. In this study, we evaluate properties of GelMA hydrogels crosslinked with visible blue light and application of these hydrogels for encapsulation of primary dorsal root ganglion (DRG) explants inside a custom microscale device. We selected low GelMA polymer concentrations of 3 % and 6 % w/v for hydrogel preparation to create soft materials and demonstrate the effect of degree of functionalization (DoF) on stiffness of GelMA in a blue light-induced crosslinking system. Additionally, a decrease in crosslinking efficiency and stiffness of GelMA was observed upon reconstitution of the polymer in DMEM cell culture media compared to PBS. To evaluate the axonal outgrowth, DRG explants were encapsulated in various GelMA formulations and cultured for 7 days. Following incubation, the explants were fixed and immunochemically labelled with anti-beta-III-tubulin antibody. We further describe an application of custom-developed semi-automated image analysis tool together with linear mixed effects modeling to evaluate axonal outgrowth across different GelMA formulations. Among the formulations tested in our study, 3 % GelMA 95 % DoF hydrogels supported the longest neurite outgrowth suggesting that matrix properties influenced neurite extension.

## 1. Introduction

Neural systems-on-a-chip are microfabricated devices that mimic the structural and functional organization of nervous system tissues and offer a versatile platform for modelling pathophysiological processes of nervous system diseases and injuries, as well as for high-throughput drug screening [1]. Such systems often integrate biomaterial scaffolds, multiple cell types, and specific topographical, mechanical and biochemical cues [1,2]. Hydrogels are biomaterials with excellent water-retention capabilities that offer 3D architecture for cell encapsulation similar to the natural extracellular matrix (ECM) [2,3]. Thus, application of hydrogels as scaffold materials in neural systems-on-a-chip devices offers an advantage to study cellular interactions, neurite growth and cell migration in 3D [1].

Gelatin methacrylate (GelMA) is a semi-synthetic hydrogel derived from gelatin via the modification of amine groups of lysine and hydroxyl residuals with methacrylate side groups [4–6]. This modification allows the GelMA prepolymer to undergo rapid polymerization into a hydrogel via light exposure in the presence of an appropriate photoinitiator [7]. Photopolymerization of GelMA is often carried with an application of UV-light (*λ* = 365 *nm*), and there exist many photoinitiators compatible with UV-light wavelengths, including LAP and Irgacure-2959 [6]. However, the use of alternative photoinitiators enables photopolymerization with longer-wavelength visible light, which is generally considered less cytotoxic than UV irradiation [8,9].

In addition to its efficient polymerization mechanism, GelMA offers exceptional biocompatibility and cell-adhesive properties due to the retention of gelatin amino-acid sequence and specifically cell-adhesive RGD-motifs [10]. Through changes in concentration, degree of methacrylation (degree of functionalization, DoF) of GelMA, and crosslinking conditions, hydrogels with different levels of stiffness can be produced, which affect cell survival and spreading [4,11,12]. Further, by altering GelMA stiffness, hydrogels with properties suitable for a specific cell type can be manufactured. GelMA hydrogels have been successfully applied for the encapsulation of a variety of cell types, such as mesenchymal stem cells [4,13,14], chondrocytes [8], fibroblasts [15], osteoblasts [16]. GelMA has also been shown to support the survival of microglial cells [17], Schwann cells [18], neural stem cells [19] and even human spinal cord organoids [20] upon encapsulation, making it a promising material for the development of biomaterial scaffolds and application in the organ-on-a-chip platforms, including neural systems-on-a-chip devices.

In this study, we investigate the application of GelMA hydrogels for encapsulation and outgrowth of primary dorsal root ganglion (DRG) neurons. A DRG is a cluster of cells located near the spinal cord. DRG neurons are classified as peripheral nervous system (PNS) neurons, projecting two axonal branches: one-to the periphery, collecting sensory information from skin and joints, and one that transmits the information into the spinal cord and synapses with secondary neurons of spinothalamic tract [21,22]. DRG neurons do not spontaneously regenerate after central nervous system (CNS) injuries, such as spinal cord injury (SCI), however, can be prompted to regenerate following the damage to their peripheral branch [21,23,24]. This property has been utilized to activate pro-regenerational transcription programs in neuron cell bodies and stimulate regeneration of the CNS branch in vivo [21,23]. DRGs contain distinct types of sensory neurons responding to different neurotrophins, as well as other types of cells such as fibroblasts and macrophages, as well as Schwann cells and satellite glial cells which are responsible for myelin formation and growth factor synthesis [25,26]. Together, these features make DRG explants a powerful ex vivo platform for studying neural development and regeneration, modeling pathophysiological processes, and evaluating therapeutic interventions while maintaining the complex interactions between neurons and supporting glial and non-neuronal cells.

In vitro DRGs have been used as a model cell system to assess 3D neural network formation in microfluidic chips [27], and outgrowth of neurites in 2D and 3D using various biomaterials [28–30]. ECM-derived hydrogel such as Matrigel [27], and hydrogels based on collagen [29], hyaluronic acid [30,31], laminin [29], and fibrin [32] have been used for DRG encapsulation, however, few studies have used photo-crosslinkable materials, and within these studies, only UV-light induced photopolymerization has been utilized [30,31].

Although GelMA has been successfully used to support microglia, Schwann cells, and reprogrammed neural precursors, a systematic understanding of how GelMA formulation parameters, including polymer concentration, stiffness, and degree of methacrylation, influence DRG explant growth and behavior in chip-compatible microphysiological systems remains limited. Here we describe application of a visible blue light photoinitiator system for GelMA crosslinking. We discuss GelMA hydrogel design considerations to improve 3D axonal outgrowth from DRGs providing a foundation for future studies of neural growth, injury, and regeneration in engineered in vitro systems, as well as for the development of biocompatible bioinks and scaffolds for CNS and PNS regeneration.

## 2. Methods

### 2.1. Hydrogel fabrication

First, a 15 % w/v GelMA stock was prepared by dissolving 500 mg of sterile lyophilizate of either PhotoGel 50 % DS (Cellink, VL3500000502) or PhotoGel 95 % DS (Cellink, 5208-1GM) in 3.35ml of the solvent of choice. Solvents used include Phosphate Buffer Saline (PBS, ThermoFisher Scientific, 21063029), Hank’s Balanced Salt Solution, no phenol red (HBSS, ThermoFisher Scientific, 14025092), and ‘base media’ without (BM) or with (BM-PHR) phenol red. BM contains a mixture of DMEM, high glucose, HEPES, no phenol red (ThermoFisher, 21063029), 1 % Penicillin-Streptomycin (ThermoFisher Scientific, 15140122) and 1 % GlutaMAX supplement (ThermoFisher Scientific, 35050061); BM-PHR: a mixture of DMEM, high glucose, with phenol red (ThermoFisher Scientific, 11965092), 1 % Penicillin-Streptomycin and 1 % GlutaMAX. The solution was placed on a hot plate set to 40 °C and left overnight under constant stirring until the complete dissolution of lyophilizate. Resulting stock solutions were GelMA 50DoF (GelMA with an average DoF of 50 %) or GelMA 95DoF (GelMA with an average DoF of 95 %). To prepare GelMA hydrogels of required concentration (3 % or 6 % (w/v)), an aliquot of 15 % GelMA stock solution was further diluted in the respective solvent with an addition of 0.2 mM of ruthenium-tris (2,2′-bipyridyl) dichloride (Ru, Sigma-Aldrich, 544981-1G) and 2 mM of sodium persulfate (SPS, Sigma-Aldrich, 216232-25G). GelMA Ru/SPS hydrogel formulations were protected from light and kept at 37 °C prior to crosslinking and used within 15 minutes. To prepare hydrogel samples for physical properties characterization, 300 µl of the formulation was poured into a circular mold (d = 15 mm, h = 1.7 mm) and placed under the LED light source (Mightex precision LED spotlight, *λ* = 450 *nm*, 63.7mW/cm^2^ light power density) set at 10 cm working distance. Hydrogel formulations were exposed to light for 60 s to complete the crosslinking.

### 2.2. Hydrogel characterization

#### 2.2.1. Swelling ratio and sol fraction experiments

The initial analysis of crosslinking efficiency and density of GelMA hydrogels was performed using sol fraction (SF) and equilibrium swelling ratio (SReq) measurements, following the methodology described by Lim et al. [8]. SF value reflects the percentage of the GelMA polymer that didn’t polymerize upon the application of light stimulus but remained within the sample. GelMA polymer can undergo reversible thermal crosslinking: transform into a gel form at room temperatures and below and transform back into a sol form at body temperature (37 °C). This property is being utilized in the sol fraction methodology.

Immediately after the crosslinking, GelMA hydrogels were transferred into the individual containers, protected from light, and placed in the –-80 °C freezer for several hours. Frozen hydrogels were lyophilized overnight, then, their initial dry mass was recorded (*m_initial_*_,*dry*_). Dried hydrogels were then submerged in 2 ml of PBS and incubated overnight at 37°C, protected from light to dissolve all uncrosslinked macromers, leaving only the photopolymerized portion behind. The next day, PBS was quickly removed, and hydrogels underwent the freeze-drying cycle again. Final dry mass was recorded as *m_dry_*. Sol fraction, % (SF) was calculated using the equation:

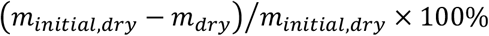

Lim et al. have previously reported differences in sol fraction of GelMA hydrogels prepared with different concentrations of Ru/SPS and varying light exposures [8]. They have reported that Ru and SPS concentrations of 0.2 and 2 mM respectively achieved “complete crosslinking” of 10 wt % GelMA during the exposure of the formulation to visible light for >30 s at 30mW/cm^2^ with SF values in the range of 15-30 % [8]. Based on this finding, 0.2/2 mM concentration of Ru/SPS was selected as a constant in our work to further explore crosslinking of lower GelMA concentrations using visible light.

For swelling ratio measurements, GelMA hydrogels were immersed in PBS immediately after the crosslinking and incubated overnight at 37 °C protected from light. The next day, hydrogels were removed from the swelling media, weighed, and their wet mass was recorded (*m_s_*_+*ollen*_). Hydrogels were placed in the –80 °C freezer for several hours followed by lyophilizing overnight. The following day, dried hydrogel mass was recorded as *m_dry_*. Equilibrium swelling ratio (SReq) was calculated using the equation:

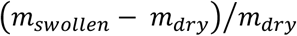

#### 2.2.2. Rheology

GelMA hydrogels were photo-crosslinked as described above inside a circular polydimethylsiloxane (PDMS) mold (diameter: 25 mm, height: 1.7 mm; diameter/height ratio within 10-50 as per ISO 6721-10). Following fabrication, samples were placed in PBS and stored at 37 °C, protected from light for 24 hours prior to analysis. Samples were analyzed in a swollen state using a 25 mm smooth parallel plate system with 0.03 N normal force control to prevent sample slippage (TA Discovery Hybrid HR 30 Rheometer). A solvent trap covered the geometry to minimize sample evaporation during measurements conducted at physiological temperature (37 °C). For viscoelastic assessment, distinct hydrogel samples were used for each of two analyses: i) oscillatory amplitude sweep (0.1-10 % at 5 rad/s) to determine the linear viscoelastic region (LVER), and ii) oscillatory frequency sweep (0.1–10 rad/s) at a strain within the identified LVER. All experimental conditions for both analyses were tested in triplicate (n=3 per test type) unless stated otherwise.

#### 2.2.3. Statistical analysis, hydrogel properties

Graphing and statistical analyses were performed using GraphPad Prism Version 10.6.1 (GraphPad Software, San Diego, CA, USA) unless stated otherwise. For comparisons between multiple groups one-way and two-way analysis of variance (ANOVA) and multiple comparisons tests with Tukey’s post-hoc test were performed. p<0.05 was considered to indicate statistical significance.

### 2.3. *In vitro* experiments

#### 2.3.1. Dorsal root ganglion explant harvest

Cell culture experiments were carried out using dorsal root ganglions explants (DRGs) freshly isolated from day 5-9 Thy1-GFP postnatal rats. Briefly, day 5-9 rat pups were sacrificed with isoflurane anesthesia followed by decapitation. A dorsal incision was made through the vertebral column from cervical to sacral. A laminectomy was performed along the spine, and the spinal cord was removed to expose DRGs. DRGs were excised from their pockets and nerve roots were trimmed prior to collection into a petri dish containing Hanks’ Balanced Salt Solution (HBSS, H9349-100ML, Sigma Aldrich). For downstream experiments, DRGs were cultured for 7 days in a basal medium consisting of Dulbecco’s Modified Eagle Medium (DMEM, ThermoFisher Scientific, 11965092), supplemented with 10 % fetal bovine serum (FBS, ThermoFisher Scientific, 12483020), 1 % penicillin-streptomycin (P/S, ThermoFisher Scientific, 15140122), 1 % GlutaMAX (ThermoFisher Scientific, 35050061), 1 x B27 (Gibco, A3582801), 100 ng/mL nerve growth factor (rNGF, CedarLane, 556-NG-100) and 10 µM cytosine β-D-arabinofuranoside hydrochloride (Ara-C, Sigma Aldrich, C6645-100MG), at 37 °C and 5 % CO_2_. DRG explant harvest was carried out in accordance with the protocol #A25-0169 approved by the UBC Animal Care Committee.

#### 2.3.2. Fabrication of PDMS microdevices for 3D cell culture

PDMS-glass microdevices were fabricated through replica molding and assembled via oxygen plasma treatment. The microdevice contains four parallel channels. Each channel contains two microwells (2 mm in diameter) connected by a central microchannel (1 mm x 1 mm x 1.5 mm). Above the microchannels is a media reservoir (1 mL). To create the PDMS layers, liquid PDMS (Sylgard 184, Dow Corning) mixed at a 10:1 (elastomer base: curing agent) ratio was poured into resin molds created using 3D printing (MiiCraft). To obtain uniform thickness a transparency film lid was placed over the PDMS-filled molds after degassing and prior to curing (overnight at 65 °C) as previously described [33]. A glass coverslip is plasma bonded to the PDMS layer to enclose the microchannels and facilitate microscopy.

Prior to use, microdevices were autoclaved and then coated with poly-D-Lysine (PDL) solution (Gibco, A3890401). Briefly, PDL stock was dissolved in UltraPure Distilled Water (ThermoFisher Scientific, 10977035) to create a 0.05 mg/ml solution. 500-800 µL of the PDL solution was added to each device to completely cover the well and channels surface area and incubated for 1-3 hours. Then, the devices were washed with UltraPure Distilled Water three times and left in the biosafety cabinet to allow for the residual evaporation. Pre-coated devices were stored at 4 °C for up to 2 weeks prior to use.

**Figure 1.**
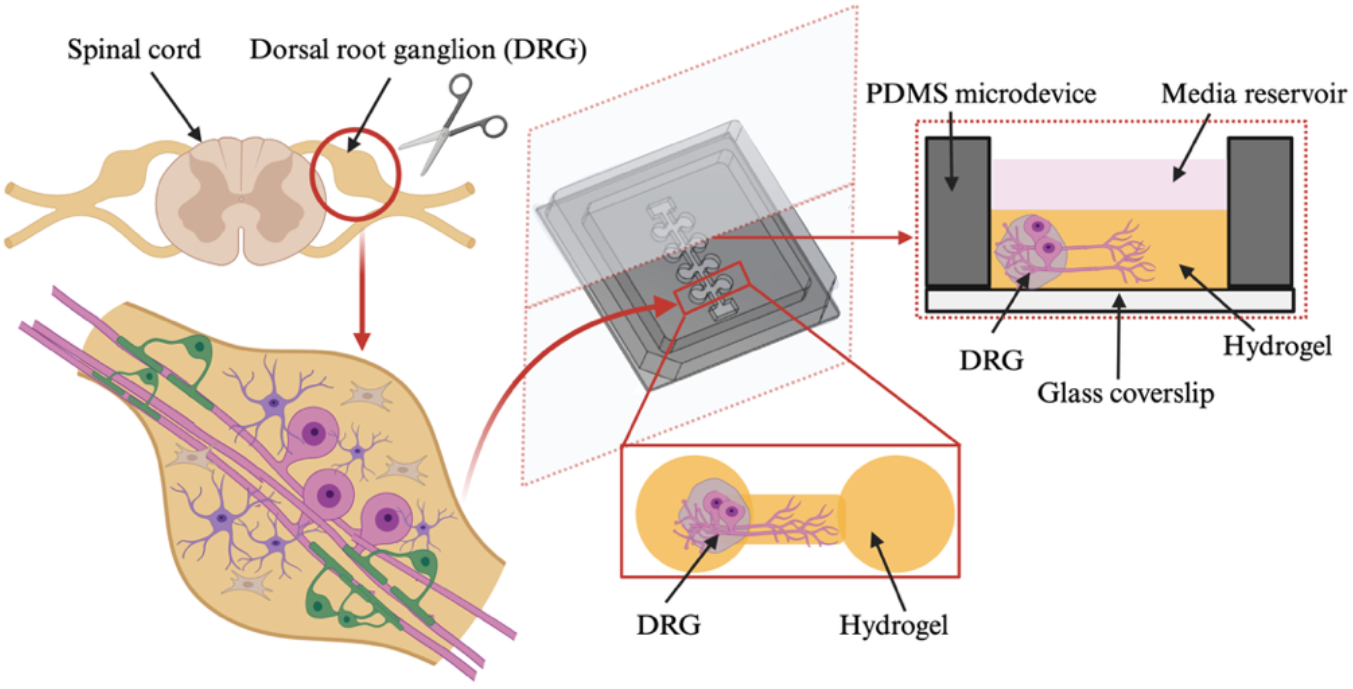
Overview of the workflow, demonstrating DRG harvest, placement of the explant into the PDMS microdevice, and a top view and a cross-sectional schematic of the device showing the position of the DRG in cell culture. Created in BioRender.

#### 2.3.3. DRG encapsulation in GelMA hydrogels

DRGs were transferred into the sterile pre-coated PDMS microdevices and each explant was placed into an individual well of the device. The wells and channels were then immediately filled with 50-70 µl of fresh GelMA Ru/SPS formulation, and the device was placed under the lens of the light source for photo-crosslinking. The lens-to-device distance was set at 10 cm to power density 63.7 mW/cm^2^ for 60 s. Immediately following the photopolymerization, the device containing DRGs encapsulated in GelMA hydrogel was filled with 1.2 mL of cell culture media and placed into the incubator (37 °C, 5 % CO2). Each PDMS microdevice contained four DRG explants encapsulated in a given GelMA hydrogel formulation.

For the positive control group, DRGs were encapsulated in the Cultrex UltiMatrix Reduced Growth Factor Basement Membrane Extract hydrogel (Ultimatrix RGF BME, R&D Systems, # BME001-05). Following the placement of explants into the wells of PDMS device, Ultimatrix was diluted in BM to 8 mg/mL concentration and used to fill out the wells and the channel of the device. The device was placed into the incubator for 20 minutes to allow for Ultimatrix crosslinking at 37 °C, then 1.2 mL of cell culture media was added on top.

#### 2.3.4. DRG explant fixation, staining, and imaging

Following the encapsulation of DRGs in hydrogels and culturing over 7 days, explants were fixed with 2 % or 4 % paraformaldehyde (PFA, in PBS), for 15 minutes at room temperature, and permeabilized with 0.3 % Triton X-100 for 15 minutes at room temperature. Following permeabilization, explants were incubated with a blocking solution containing 2 % bovine serum albumin (BSA), 2 % normal goat serum, and 0.1 % Tween-20 in PBS for 1 hour at room temperature. Explants were then incubated with primary antibody anti-β-Tubulin III clone TUJ1 (1:1000, STEMCELL Technologies, 60052) in blocking solution for 48 hours at 4 °C. Explants were then incubated with goat anti-mouse IgG (H+L) cross-adsorbed secondary antibody (Alexa Fluor™ 647, 1:500, Abcam, A21235) in blocking solution for 24 hours at 4 °C. Finally, explants were incubated with nucleic acid stain Hoechst (1:2000 in PBS, ThermoFisher Scientific, 66249) for 10 minutes at room temperature and stored in PBS until imaging.

Images were acquired using a Leica THUNDER Imager at 5 x magnification. Widefield fluorescence 3D Z-stacks were obtained at 15 µm intervals in the bright field and fluorescent (AF688) channels followed by Large Volume Computational Clearing (LVCC) as the clearing method. Computational Clearing was applied to reduce out-of-focus background fluorescence while preserving in-focus signal, improving contrast and visualization of structures throughout the imaging volume.

12 different GelMA formulations (conditions) were applied for DRG explant encapsulation, and the outgrowth of neurites from DRGs in these hydrogels was analyzed from collected images using the custom-developed Automated Quantification Software tool described below.

### 2.4. Image analysis

#### 2.4.1. Explant Image Pre-Selection

DRG explants were inspected under bright field microscopy prior to image acquisition. Explants with no neurite extensions visible under bright field imaging were not imaged further. These rejected explants were manually annotated in the data set as experimental conditions that had been performed but not imaged. For those explants which were imaged, an additional manual preview operation was performed to further reject samples with insufficient image contrast or significant imaging artefacts, and to verify data formatting prior to using the semi-automated batch processing quantification software. This manual preview operation allowed for annotation of the images which had imaging artifacts (non-neuronal fluorescent debris). These images were therefore selected for further intervention during the semi-automated batch processing to manually remove artifacts that otherwise could be incorrectly segmented as neurite structures.

#### 2.4.2. Semi-Automated Quantification Software

A custom Python program was written to automate the segmentation and quantification of neurites within the DRG images. This program utilized the open source readlif library to read and work with the LIF file format provided by the Leica THUNDER Imager, and the OpenCV library for foundational image manipulation functions. Using the captured bright-field image of the explant (Figure S 1(a)), the approximate centroid and an enclosing contour of the explant core were determined and used to generate a binary mask. A second binary mask was generated from the bright-field image corresponding to the total internal region of the PDMS device in which the explants were cultured. The union of these two binary masks was used to isolate the interior region of the PDMS device in which neurites are considered possible to grow (Figure S 1(a)).

The image in the fluorescent channel with neurites labelled with β-tubulin was captured in the form of a Z-Stack (Figure S 1(b)). The quantification software processed each layer of this stack in sequence. First, the layer was binarized using a local thresholding algorithm to account for the observed spatial variation in background intensity across layers and images. Next, the previously described combined binary mask was applied to remove signals from regions of the device in which neurite growth is not considered possible. Then, the layer was processed using connected component analysis to reject disconnected speckles below a given size threshold. Additionally, this operation was used to reject regions with both high circularity and high compactness, thereby excluding small, rounded debris or other non-neurite structures from the analysis. Finally, using the estimated location of the DRG centroid, the distance to each non-zero pixel was computed. The distribution of these distances was then computed. This distribution was then inspected for gaps of a minimum size where no candidate pixels were identified. If such a gap were present, all candidate pixels with larger distances were rejected as having no connection to the centroid. Each segmented layer was re-combined into a Z-Stack for further analysis.

The segmented Z-Stack was then flattened to a flat binary image to prevent multi-counting the same neurite during quantification (Figure S 1(d)). If an image was annotated as requiring manual intervention during the preview step, the flattened image was displayed, and incorrectly segmented regions were manually removed before quantification. Quantification consisted of a simplified implementation of Sholl analysis. Using the flattened binary image of neurite pixels, the distance of each non-zero pixel to the DRG centroid was calculated. The distribution of these distances was calculated, and the distribution and median distance were reported (Figure S 1(e)). The experimental conditions which were annotated as not being imaged were assigned a median length value of 0.

#### 2.4.3. Statistical analyses, explants and neurites

First, an Exploratory Data Analysis (EDA) was performed to combine information about the number of explants tested for each condition and experimental dates with the dataset of median neurite length values obtained from images. The % of zero-growth data points (the absence of extension growth) was calculated for each condition as % of zero-growth = (number of explants with the absence of extensions/total number of explants for GelMA condition X) *100 %. From EDA, descriptive statistics for each tested condition were obtained and initial evaluation of the effect of different GelMA formulations on % of zero-growth and median neurite lengths values was performed.

However, due to the unbalanced experimental design, GelMA formulations and their impact on neurite outgrowth were further evaluated by fitting a linear mixed-effects model (LMM). This model accounts for different sources of variability by separating the variables that must be quantified and interpreted (called fixed effects), such as hydrogel formulation, and variables not needed for interpretation which are still important to consider (called random effects), such as sources of biological variability – animals, cell passages etc.

Fitting of LMM was done in Matlab using function *fitlme*. GelMA formulation parameters, such as GelMA concentration, GelMA DoF, and GelMA solvent were specified as fixed effects - variables of interest being controlled in the experiment which are contributing to the outgrowth of neurites. Animal litter (specified as experimental date) used for DRG harvest and GelMA stock (specified as a serial number of PhotoGel vial used to prepare the 15 % stock) were specified as random effects – parameters contributing to the variability in neurite outgrowth but not directly controlled in the experimental design. ‘First Model’ was applied to 174 observations (median neurite lengths data points equal to total n of explants) pulled from the dataset of 6 GelMA formulation groups specified in Table 1. ‘Second Model’ was applied to selected GelMA formulations listed in Table 2 which were specified as a fixed effect, whereas animal litter was specified as random effect. This model contained 145 observations.

**Table 1.** Experimental conditions and corresponding sample sizes (DRG explants, PDMS microdevices, and animal litters) for the GelMA formulations included in the first linear mixed-effects model.

| Condition | Sample name | n of explants | n of PDMS microdevices | n of animal litters |
| --- | --- | --- | --- | --- |
| 3% GelMA 50%DoF PBS | G3-50DoF-PBS | 18 | 4 | 4 |
| 3% GelMA 50%DoF BM-PHR | G3-50DoF-BM-PHR | 38 | 8 | 8 |
| 3% GelMA 95%DoF PBS | G3-95DoF-PBS | 25 | 6 | 6 |
| 3% GelMA 95%DoF BM-PHR | G3-95DoF-BM-PHR | 42 | 9 | 9 |
| 6% GelMA 50%DoF PBS | G6-50DoF-PBS | 30 | 6 | 6 |
| 6% GelMA 95%DoF PBS | G6-95DoF-PBS | 21 | 4 | 4 |

**Table 2.** Experimental conditions and corresponding sample sizes (DRG explants, PDMS microdevices, and animal litters) for the GelMA formulations included in the second linear mixed-effects model.

| Condition | Sample name | n of explants | n of PDMS microdevices | n of animal litters |
| --- | --- | --- | --- | --- |
| 3% GelMA 95%DoF PBS | G3-95DoF-PBS | 25 | 6 | 6 |
| 3% GelMA 95%DoF BM-PHR | G3-95DoF-BM-PHR | 42 | 9 | 9 |
| 3% GelMA 95%DoF HBSS | G3-95DoF-HBSS | 16 | 3 | 3 |
| 3% GelMA 95%DoF BM+10mM GSH | G3-95DoF-BM+10mMGSH | 21 | 6 | 5 |
| Ultimatrix | Ultimatrix | 41 | 11 | 11 |

Further, the relationship between neurite outgrowth and GelMA hydrogel stiffness was assessed through simple linear regression in GraphPad Prism (Version 10.6.1). Mean neurite length values were paired with corresponding hydrogel storage modulus (G’) values.

## 3. Results

### 3.1. GelMA crosslinking efficiency and density

There was a significant increase in the SF for the 3 % hydrogels prepared using the BM compared to PBS or BM-PHR (60.17±33.3 %, 16.75±8.67 % and 25.81±5.6 % respectively), indicating a substantial loss of mass during the 24 h incubation for BM (Figure 2 (a, b)). No difference in SF was observed between the hydrogels prepared in PBS and BM-PHR. As expected, there was a decrease in the SF values for 6 % GelMA hydrogels compared to 3 %, however, substantial difference was only observed for the hydrogels prepared using BM, but not PBS or BM-PHR. For 50DoF GelMA hydrogels prepared in PBS using 0.2/2 mM of Ru/SPS and polymerized using 60s of visible blue light exposure, SF values were 7.95±2.73 and 16.75±8.68 % for 6 % and 3 % GelMA respectively. A similar trend was observed for 95DoF GelMA with SF values of 10.45±4.73 and 13.93±1.17 for 6 % and 3 % GelMA hydrogels, respectively. Interestingly, difference in SF between the hydrogels prepared using 50DoF GelMA and 95DoF GelMA was negligible, regardless of the solvent or GelMA concentration used. 3 % GelMA-BM-PHR hydrogels have demonstrated significantly lower SF values (25.81±5.6 % and 26.56±7.15 % for 50DoF and 95DoF respectively) compared to BM, but still higher compared to PBS.

**Figure 2.**
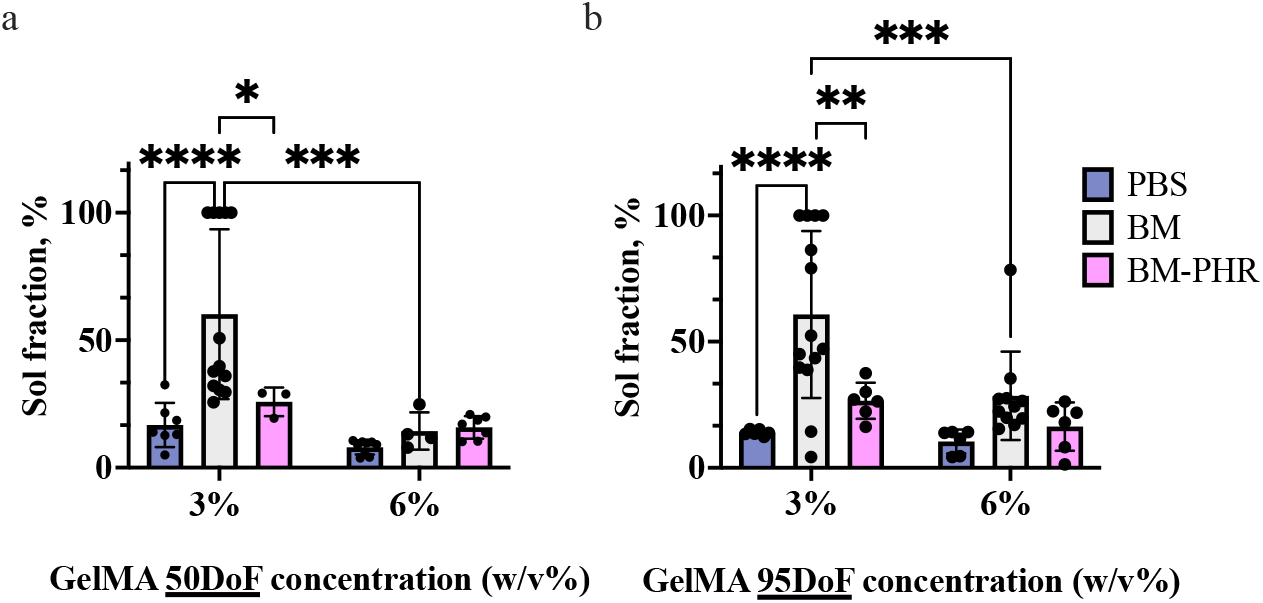
Sol fraction of hydrogels prepared with 50DoF (a) and 95DoF (b) GelMA at 3 % and 6 % (w/v) using PBS, BM, or BM-PHR. Error bars represent mean ± SD; individual data points represent technical replicates. Statistical significance is indicated by asterisks (*p<0.05, **p<0.01, ***p<0.005, ****p<0.001).

As expected, there was a significant increase in the SReq in 3 % GelMA hydrogels prepared with PBS compared to 6 % GelMA (24.39±4.52 and 19.43±1.73 for 50 % DoF, and 24.39±4.52 and 25.16±1.74 for 95DoF respectively) (Figure 3 (a)). Preparation of 6 % GelMA hydrogel in BM has resulted in a substantial increase in the SReq (35.6±6.08 and 29.7±4.26 for 50DoF and 95DoF respectively) compared to the same formulations in PBS (Figure 3 (b)). There was also a consistent increase in SReq values for 6 % GelMA 50DoF compared to 95DoF prepared in either PBS or BM.

**Figure 3.**
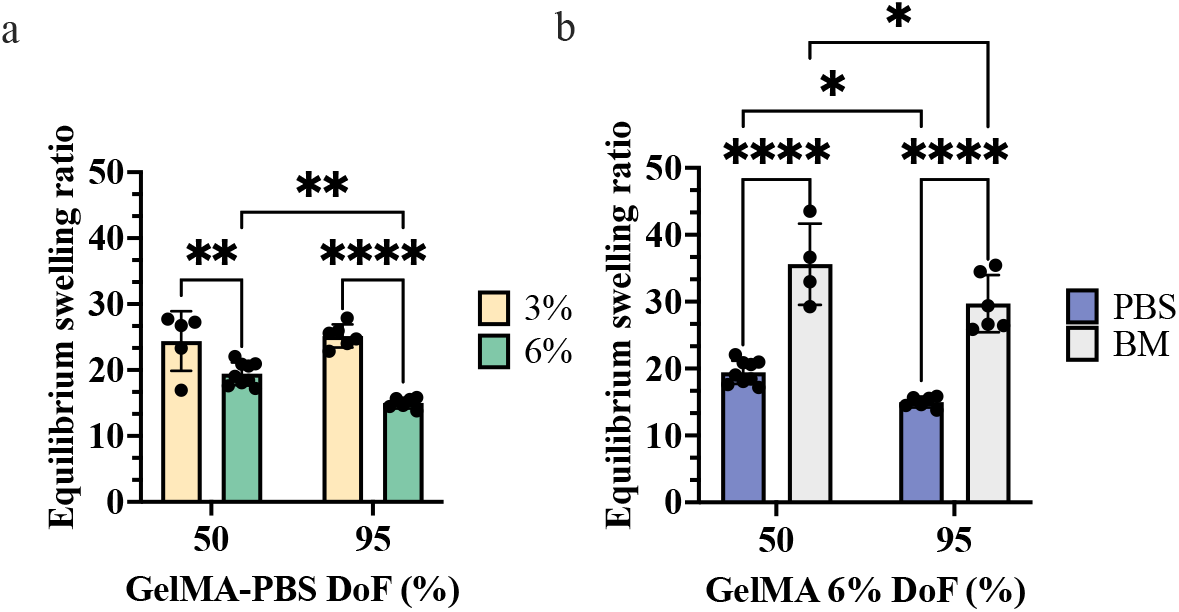
Equilibrium swelling ratio of hydrogels prepared with (a) 50DoF or 95DoF GelMA-PBS at 3 % or 6 % (w/v), and (b) 6 % GelMA prepared in PBS or BM. Hydrogel samples were swollen in PBS for 24 h. Error bars represent mean ± SD; individual data points represent technical replicates. Statistical significance is indicated by asterisks (*p<0.05, **p<0.01, ***p<0.005, ****p<0.001).

Interestingly, the addition of 5 mM GSH to the BM, resulted in a significant decrease in SF for 3 % GelMA 95DoF hydrogels, with a further decrease observed as the GSH concentration increased to 10 mM (31.04 ± 2.91 % and 4.97 ± 2.7 % for 5 mM and 10 mM GSH, respectively) (Figure 4 (a)). A similar trend was observed for the 3 % GelMA 50DoF hydrogels, however, the decrease in SF was only noticeable for the addition of 10 mM GSH to BM, but not 5 mM (30.61 ± 3.63 % and 21.7 ± 7.53 % for 5 mM and 10 mM GSH, respectively) (Figure 4 (b)). Notably, for 3 % GelMA hydrogels prepared in PBS or BM-PHR, no significant differences in SF were observed upon the addition of GSH. (Figure 4 (c,d)).

**Figure 4.**
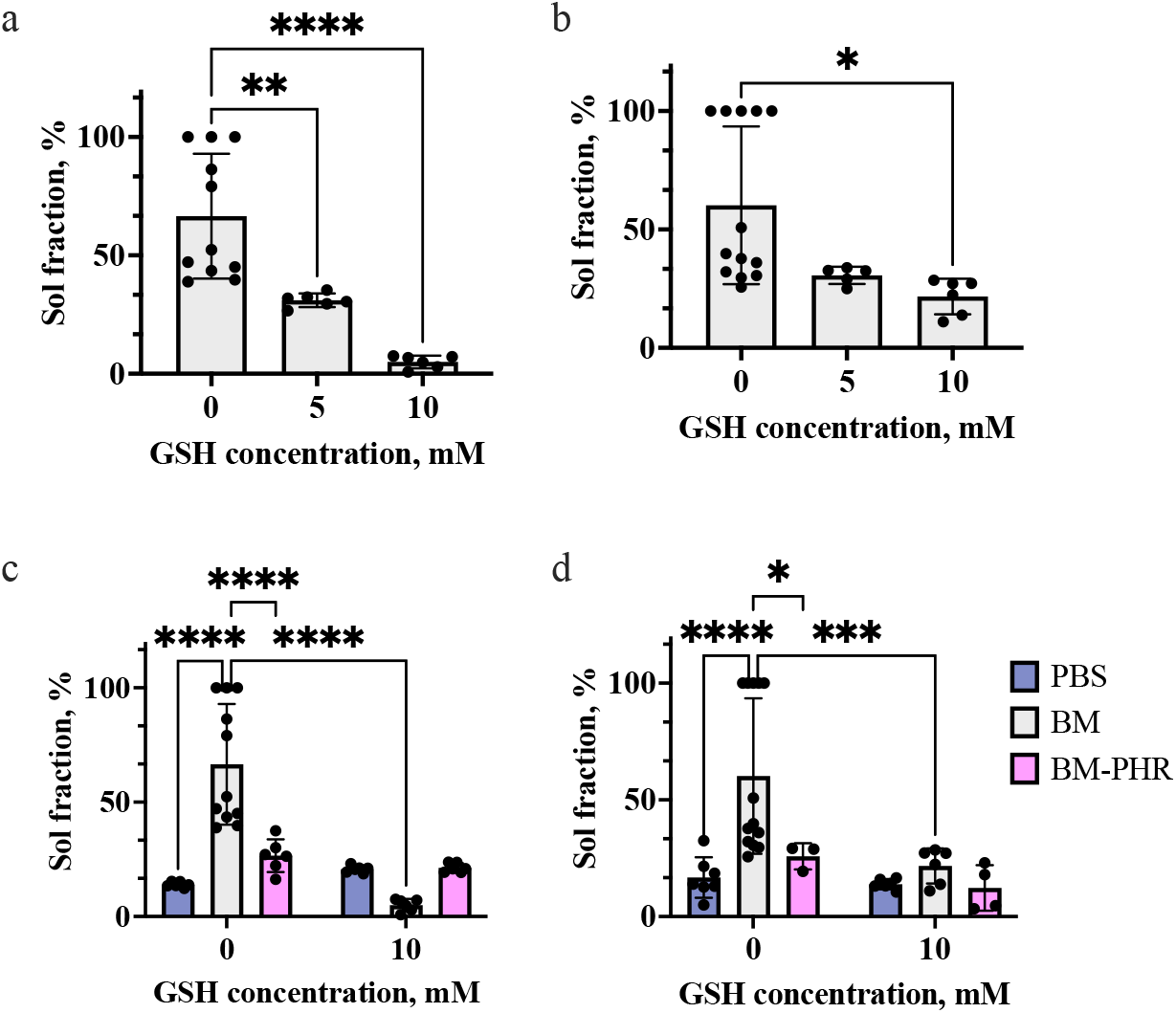
Sol fraction of 3 % GelMA 95DoF (a) or 50DoF(b) hydrogels prepared in BM with 0,5 or 10mM of GSH, and 3 % GelMA 95DoF (c) or 50DoF (d) hydrogels prepared) using PBS, BM, or BM-PHR in the absence or with an addition of 10 mM of GSH. Error bars represent mean ± SD; individual data points represent technical replicates. Statistical significance is indicated by asterisks (*p<0.05, **p<0.01, ***p<0.005, ****p<0.001).

### 3.2. GelMA hydrogel stiffness

The storage moduli (G’) of GelMA hydrogels were analyzed once samples reached equilibrium swelling in PBS. For GelMA hydrogels prepared using PBS, 3 % GelMA hydrogel demonstrated a substantial decrease in G’ compared to the 6 % hydrogels for both 50DoF and 95DoF groups (G’ = 106.63±18.81 Pa and 281.65±87.95 Pa for 3 % and 6 % GelMA 50DoF respectively) (Figure 5 (a)). Notably, GelMA 95DoF hydrogels had significantly lower G’ values compared to the 50DoF hydrogels (G’ = 42.26 ± 5.3 Pa and 124.27 ± 31.12 Pa for 3 % and 6 % GelMA 95DoF respectively).

**Figure 5.**
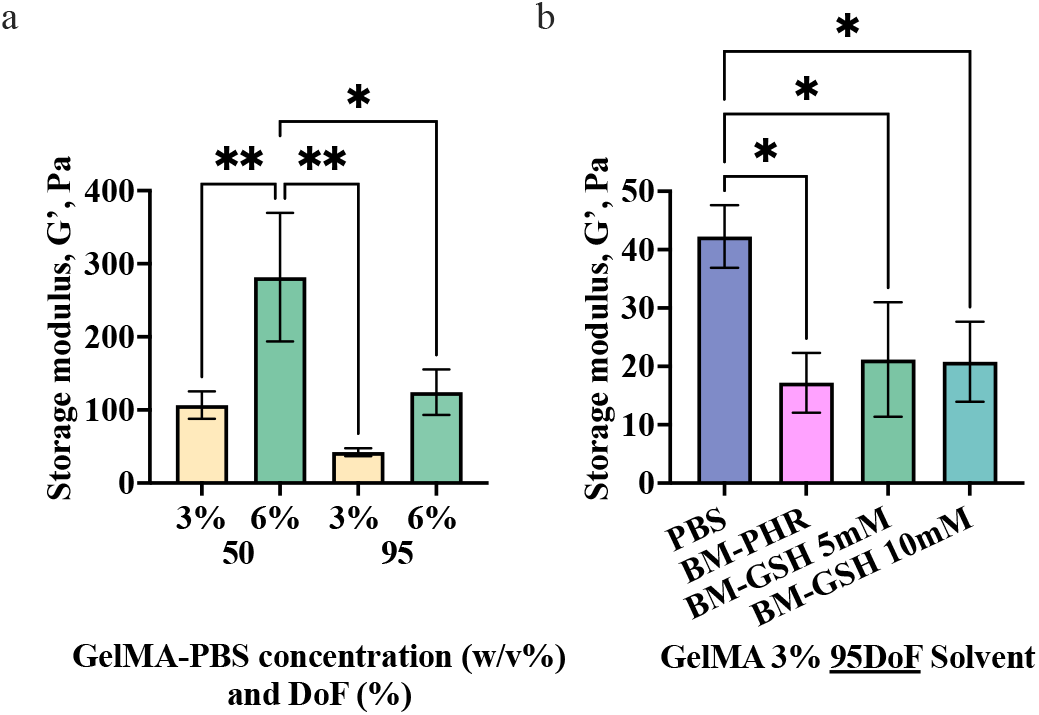
Storage moduli of 3 % and 6 % GelMA 50DoF and 95DoF reconstituted with PBS (a), and 3 % GelMA 95DoF reconstituted with PBS, BM-PHR or BM with addition of 5 or 10mM GSH. Error bars represent mean ± SD, n=3. Statistical significance is indicated by asterisks (*p<0.05, **p<0.01, ***p<0.005, ****p<0.001).

The effect of different solvents on the stiffness of hydrogels was investigated for the 3 % GelMA 95DoF formulation (Figure 5 (b)). A significant decrease in G’ value was observed when BM-PHR was used to prepare the hydrogels compared to PBS (G’ = 17.19±5.12 Pa and G’ = 42.26 ± 5.3 Pa for BM-PHR and PBS respectively). 3 % GelMA 95DoF hydrogels prepared using the BM has demonstrated very poor crosslinking and were not possible to analyze, however, upon the addition of GSH (both 5 mM and 10 mM) to BM, an increase in the G’ values was observed (G’ = 21.17 ± 9.81 Pa and 20.79 ± 6.86 Pa, respectively).

Further, the effect of light exposure on hydrogel stiffness was investigated for 6% GelMA 50DoF and 95DoF formulations (Figure 6). Samples were placed into the rheometer and analyzed immediately following light exposure, in a non-swollen state. No significant difference was observed for 6% GelMA 50DoF hydrogels crosslinked for 30, 60, 90 and 120 s. However, for 6% GelMA 95DoF there was a significant increase in storage modulus for the sample crosslinked for 60 s compared to 30 s (G’ = 406.33 ± 64.47 Pa and 175.64 ± 47.48 Pa, respectively). 6% GelMA 50DoF hydrogels in a non-swollen state also have exhibited significantly higher G’ compared to 6% GelMA 95DoF samples (G’ = 550.76 ± 95.66 Pa and 406.33 ± 64.47 Pa for 50DoF and 95DoF samples crosslinked for 60 s, respectively). Notably, samples in non-swollen state exhibited significantly higher storage moduli compared to hydrogels which have reached equilibrium swelling: 406.33 ± 64.47 Pa and 124.27 ± 31.12 Pa for 6% GelMA 95 DoF, respectively (p = 0.0073, t-test).

**Figure 6.**
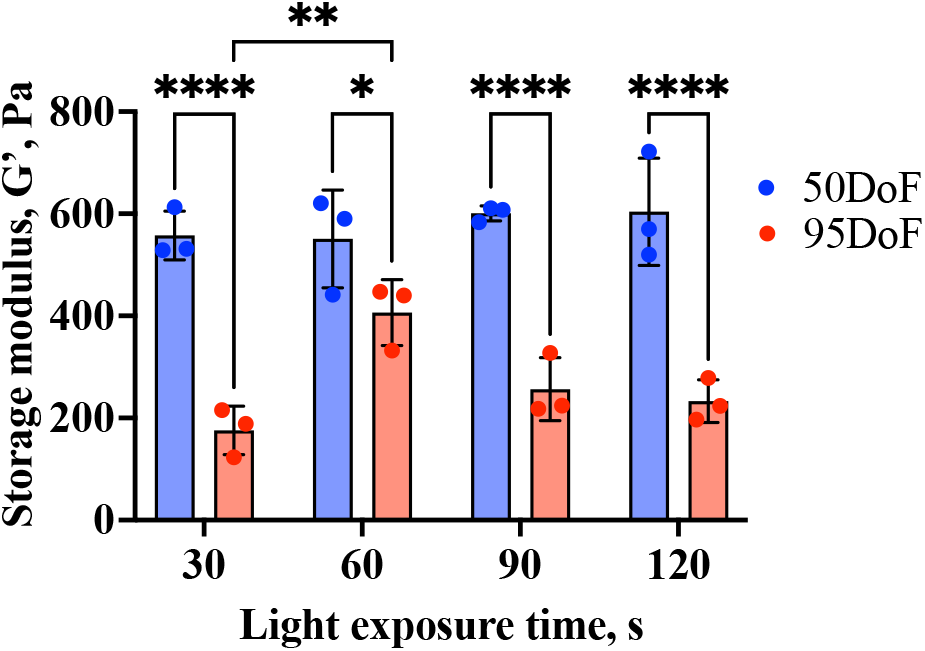
Storage moduli of 6 % GelMA 50DoF and 95DoF crosslinked through 30, 60, 90 or 120 s of blue light exposure. Error bars represent mean ± SD, n=3. Statistical significance is indicated by asterisks (*p<0.05, **p<0.01, ***p<0.005, ****p<0.001).

**Figure 7.**
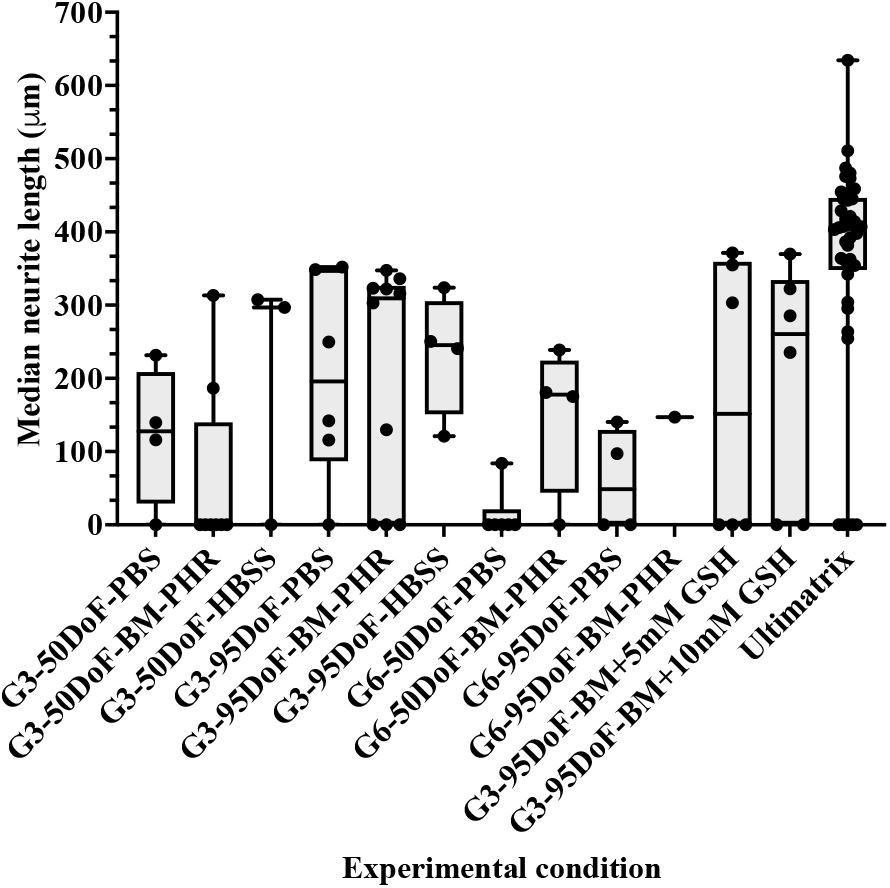
Experimental conditions included in EDA and the distribution of median neurite length values per condition. Individual values plotted are the median neurite length values per animal litter tested.

### 3.3. DRG neurite outgrowth in GelMA hydrogels

The summary of the data collected from the images of DRG explants in various GelMA hydrogels is presented in Table 3. A different number of DRG explants have been tested with each GelMA condition, creating an unbalanced experimental design. First, for each GelMA formulation, we determined the percentage of DRG explants that did not exhibit detectable neurite outgrowth after 7 days in culture. These explants were classified as “zero-growth” explants. The absence of neurite outgrowth provides a complimentary measure of the ability of each hydrogel formulation to support the initiation of neurite outgrowth which can be affected by dense hydrogel network or photocrosslinking conditions that affect neuronal viability and extension growth. This metric complements subsequent analyses of neurite extension among explants that exhibited growth which provides an insight if the formulation provides enough matrix cues for axons to extend throughout the hydrogel.

**Table 3.** Experimental conditions with corresponding sample sizes (n of explants, n of litters and n of PDMS microdevices), and EDA summary demonstrating the number of explants with the absence of growth (absolute value and percentage), as well as median and mean neurite length values per condition delivered by the image analysis tool.

| | Experimental condition | n of explants | n of PDMS microdevices | n of animal litters | n of zero-growth data points | % of zero-growth | Median neurite length, $\mu\text{m}$ | Mean neurite length, $\mu\text{m}$ |
| --- | --- | --- | --- | --- | --- | --- | --- | --- |
| 1 | G3-50DoF-PBS | 18 | 4 | 4 | 11 | 61.11 | 0 | 117 |
| 2 | G3-50DoF-BM-PHR | 38 | 8 | 8 | 25 | 65.79 | 0 | 111.5 |
| 3 | G3-50DoF-HBBS | 16 | 3 | 3 | 6 | **37.50** | 260.3 | 191.1 |
| 4 | G3-95DoF-PBS | 25 | 6 | 6 | 6 | **24.00** | 283.8 | 211.2 |
| 5 | G3-95DoF-BM-PHR | 42 | 9 | 9 | 10 | **23.81** | 286.9 | 228.2 |
| 6 | G3-95DoF-HBBS | 16 | 4 | 4 | 4 | **25.00** | 245.4 | 225.7 |
| 7 | G6-50DoF-PBS | 30 | 6 | 6 | 26 | 86.67 | 0 | 35.22 |
| 8 | G6-50DoF-BM-PHR | 14 | 4 | 4 | 8 | 57.14 | 0 | 116.3 |
| 9 | G6-95DoF-PBS | 21 | 4 | 4 | 16 | 76.19 | 0 | 72.97 |
| 10 | G6-95DoF-BM-PHR | 4 | 1 | 1 | 2 | 50.00 | 147.1 | 201.8 |
| 11 | G3-95DoF-BM+5mMGSH | 21 | 6 | 5 | 9 | **42.86** | 206.6 | 186.8 |
| 12 | G3-95DoF-BM+10mMGSH | 21 | 6 | 5 | 9 | **42.86** | 235.4 | 176.5 |
| 13 | Ultimatrix | 41 | 11 | 11 | 5 | **12.20** | 407.2 | 360.8 |

Six formulations have demonstrated <50 % explants with zero-growth and all these formulations contained a low (3 % w/v) concentration of GelMA. Out of these six formulations, five were GelMA 95DoF. The three formulations that presented with the lowest percent of zero-growth (<30 %) were 3 % GelMA 95DoF prepared with different solvents (HBSS, PBS and BM-PHR). 3 % GelMA 95DoF prepared in HBSS, PBS, or BM-PHR have also demonstrated the highest median neurite lengths: 260.3, 283.8 and 286.9 µm respectively. 3 % GelMA 50DoF or 95DoF in HBSS also demonstrated high median neurite length values of 245.4 and 260.3 µm, respectively. Interestingly, 3 % GelMA 50DoF prepared in PBS or BM-PHR demonstrated high percent of zero-growth (>60 %) and low median neurite length values. 3 % GelMA 95DoF prepared in BM+GSH demonstrated moderate percent of zero-growth (43 %) but high median neurite length values of 206.6 and 235.4 µm for 5 mM and 10 mM GSH, respectively. All hydrogels prepared with 6 % GelMA demonstrated high percent of zero-growth (>50 %) and low median and mean neurite lengths values (<200 µm). The positive control hydrogel Ultimatrix has demonstrated the lowest percent of zero-growth (12.2 %) and the highest median neurite length value of 407.2 µm.

In Table 3, median neurite length values for each experimental day are plotted per GelMA condition. The scattered nature of the values indicates the variability in neurite outgrowth observed on each experimental day with different animal litters. On some experimental days, for 3 % GelMA 95DoF conditions (in HBSS, PBS, BM-PHR and BM+10mM GSH) the majority of median neurite length values have clustered above 200 µm, however, there are also values at 0 µm for other experimental days. Further, we have observed high variability in neurite outgrowth between individual explants encapsulated in the same hydrogel formulation within one device. Figure 8 illustrates the variability in neurite outgrowth among individual DRG explants and across animal litters for two representative GelMA formulations. For both formulations, substantial variability in median neurite length was observed among individual explants within the same experiment, as well as differences in the distribution of neurite lengths across litters. Individual explants with no detectable neurite outgrowth were also observed. This biological heterogeneity motivated the use of a linear mixed-effects model to evaluate the effects of GelMA formulation while accounting for potential variability associated with animal litter.

**Figure 8.**
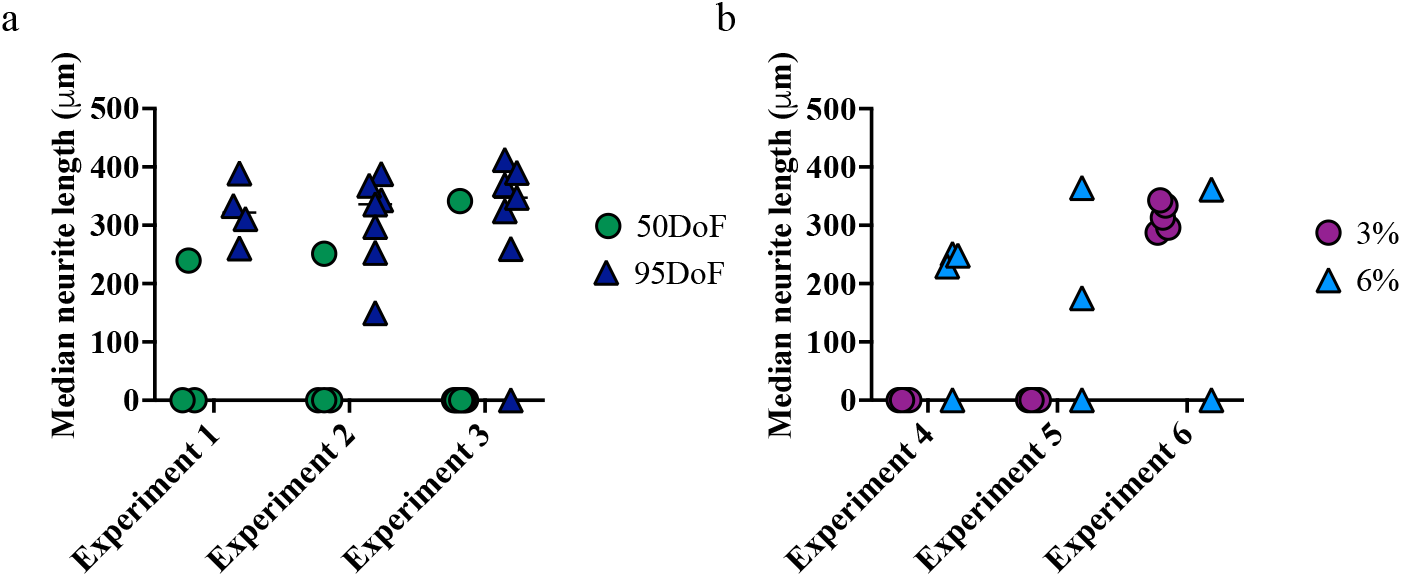
Distribution of median neurite length data points for every explant tested on different experimental days with different animal litters with (a) 3 % GelMA – BM-PHR 50DoF or 95DoF, and (b) GelMA 50DoF BM-PHR 3 % or 6 % (w/v).

The first LMM investigated the combinations of 3 % or 6 % GelMA, 50DoF or 95DoF, PBS or BM-PHR solvents used for hydrogel fabrication and their effect on neurite length. 6 % GelMA and 50DoF in PBS were used as references to which all other comparisons were made, and the baseline average neurite length for the combination of reference GelMA parameters was estimated to be 4.39 µm. The variability between the animal litters used for DRG harvests, which was a total of 20 different litters, was estimated to contribute to the 54.43 µm difference in neurite length. Similarly, the variability between different GelMA stocks, which was a total of 7 different stocks serial numbers, was estimated to contribute to the 41.54 µm difference in neurite length. The residual error contributing to unspecified variability between the individual explants was estimated to be 131.84 µm.

After accounting for random variability, it has been estimated that decreasing GelMA concentration to 3 % has produced neurite lengths that are on average 110.24 µm longer compared to 6 % GelMA (p<0.01, 95 % CI). Further, the transition to GelMA 95DoF has produced neurite lengths that are on average 96.02 µm longer than GelMA 50DoF (p<0.05, 95 % CI). Switching from PBS to BM-PHR for hydrogel fabrication did not yield a significant difference in DRG neurite length for respective hydrogel samples. Marginal R^2^_m_ for the model was calculated according to Nakagawa et.al [34] as the ratio of variance explained by fixed effects (such as GelMA concentration, DoF and solvents) to total variance (fixed, random and residual) and was determined to be 0.212. Conditional R^2^_c_ of the model was estimated to be 0.314 representing the proportion of variance explained by both fixed and random effects relative to the total variance, meaning residual 0.685 of data variance is due to unspecified variability in neurite length between the individual explants.

Further, predicted neurite lengths were estimated from the linear mixed-effects model for each combination of GelMA concentration and degree of functionalization, the fixed effects identified as significant. These model-based predictions isolate the effects of the fixed experimental factors, while accounting for random variability across animal litters and GelMA stocks. On Figure 9 model-based predicted values (b) are plotted alongside the raw data (a). The trend of increased neurite lengths for 3 % GelMA 95DoF can be observed on plots for both model-based predicted values and the raw data serving as validation of the model.

**Figure 9.**
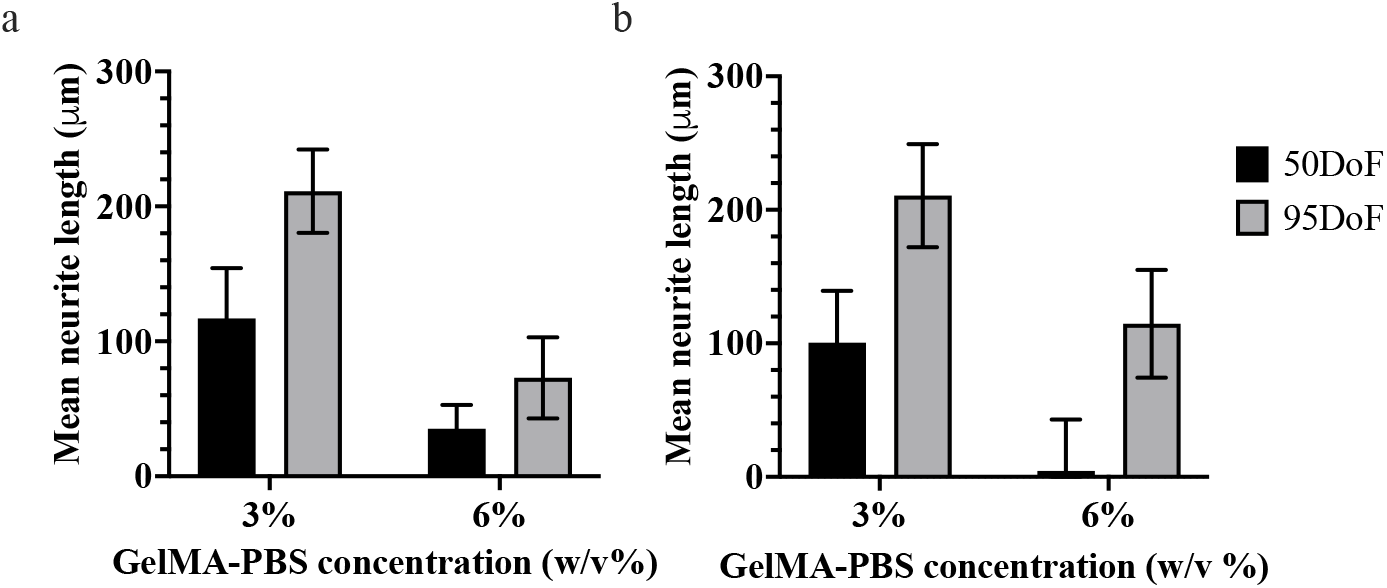
Mean neurite length data for 3 % and 6 % (w/v), 50DoF and 95DoF GelMA-PBS formulations obtained from image analysis tool (a), from the linear mixed effects model (b).

**Figure 10.**
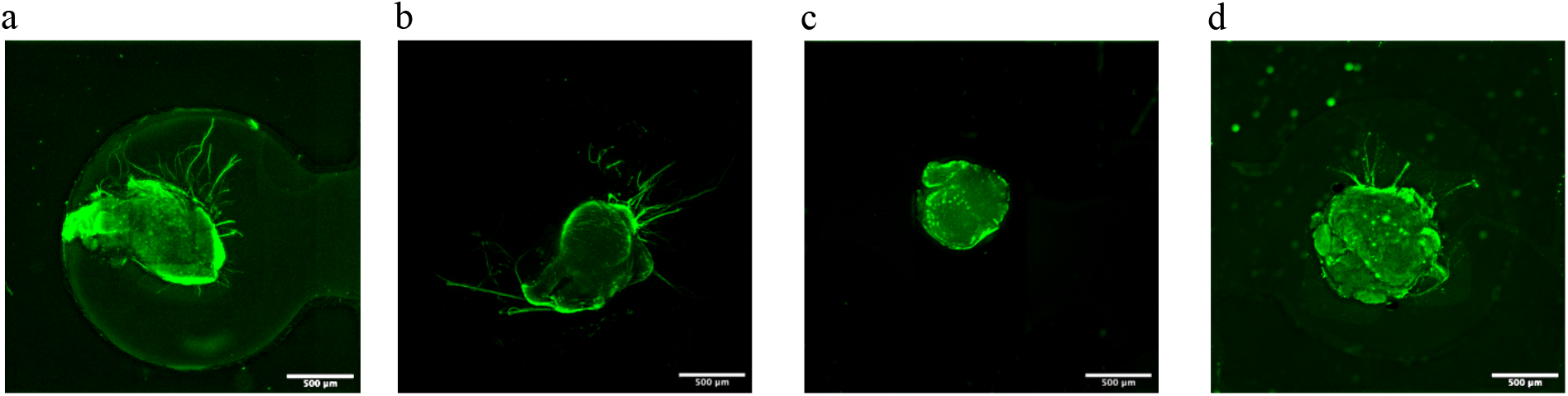
Representative images of DRGs in 3 % GelMA 50DoF (a), 95DoF (b), 6 % GelMA 50DoF (c), and 95DoF (d). Scale bar is 500μm. Brightness and contrast of the images were enhanced, and background noise was removed through ImageJ software for demonstration purposes.

Since 3 % GelMA 95DoF has demonstrated the best neurite outgrowth among the GelMA formulations tested above, we have evaluated how this formulation compares to the positive control hydrogel and other 3 % GelMA 95DoF formulations prepared with various solvents by fitting another LMM to the dataset of median neurite lengths values. 3 % GelMA 95DoF - BM-PHR was set as a reference condition with predicted average neurite length of 243.82 µm. The variability between the animal litters used for DRG harvests (total 15 different litters/experiment days) was estimated to contribute insignificantly to the <0.1 µm difference in neurite length. The residual error contributing to unspecified variability between the individual explants was estimated to be 152.77 µm.

Switching from BM-PHR to PBS, HBSS or BM+10 mM GSH for 3 % GelMA 95DoF fabrication has not demonstrated a statistically significant shift in neurite outgrowth, with these formulations exhibiting a decrease in average neurite length by 32.56, 18.09 and 67.32 µm respectively (p>0.1, 95 % CI). On the other hand, Ultimatrix was estimated to produce average neurite length 116.96um longer compared to the reference GelMA treatment (p<0.01, 95 % CI).

The marginal and conditional R^2^ values were both calculated to be 0.162, indicating that the fixed effects included in the model, such as hydrogel formulation, explained approximately 16% of the total variability in neurite length, while the specified random effects (animal litter and GelMA stock) contributed negligibly to the observed variance. The remaining unexplained variance (83.8%) likely reflects biological heterogeneity among individual DRG explants, which is expected in primary tissue cultures. Despite the residual variability, the linear mixed-effects model identified significant effects of hydrogel formulation by explicitly accounting for the hierarchical structure of the experimental data.

Model predictions were generated to demonstrate expected neurite lengths for each evaluated hydrogel formulation. On Figure 11 model-based predicted values (b) are plotted alongside the raw data (a). The trend of increased neurite lengths for positive control Ultimatrix and non-significant difference in average neurite length between four different 3 % GelMA 95DoF formulations can be observed on both plots serving as validation of the model.

**Figure 11.**
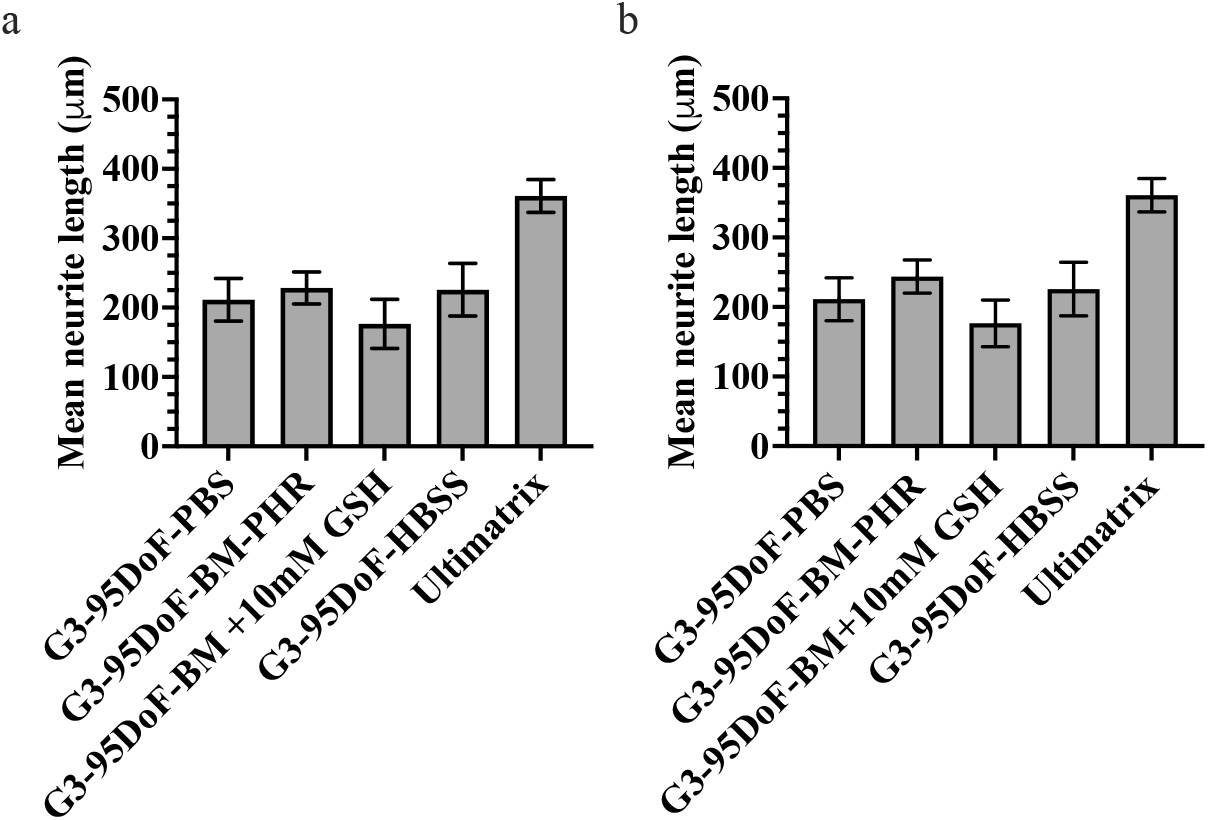
Mean neurite length data for GelMA and Ultimatrix formulations obtained from image analysis tool (a), from the linear mixed effects model (b).

**Figure 12.**
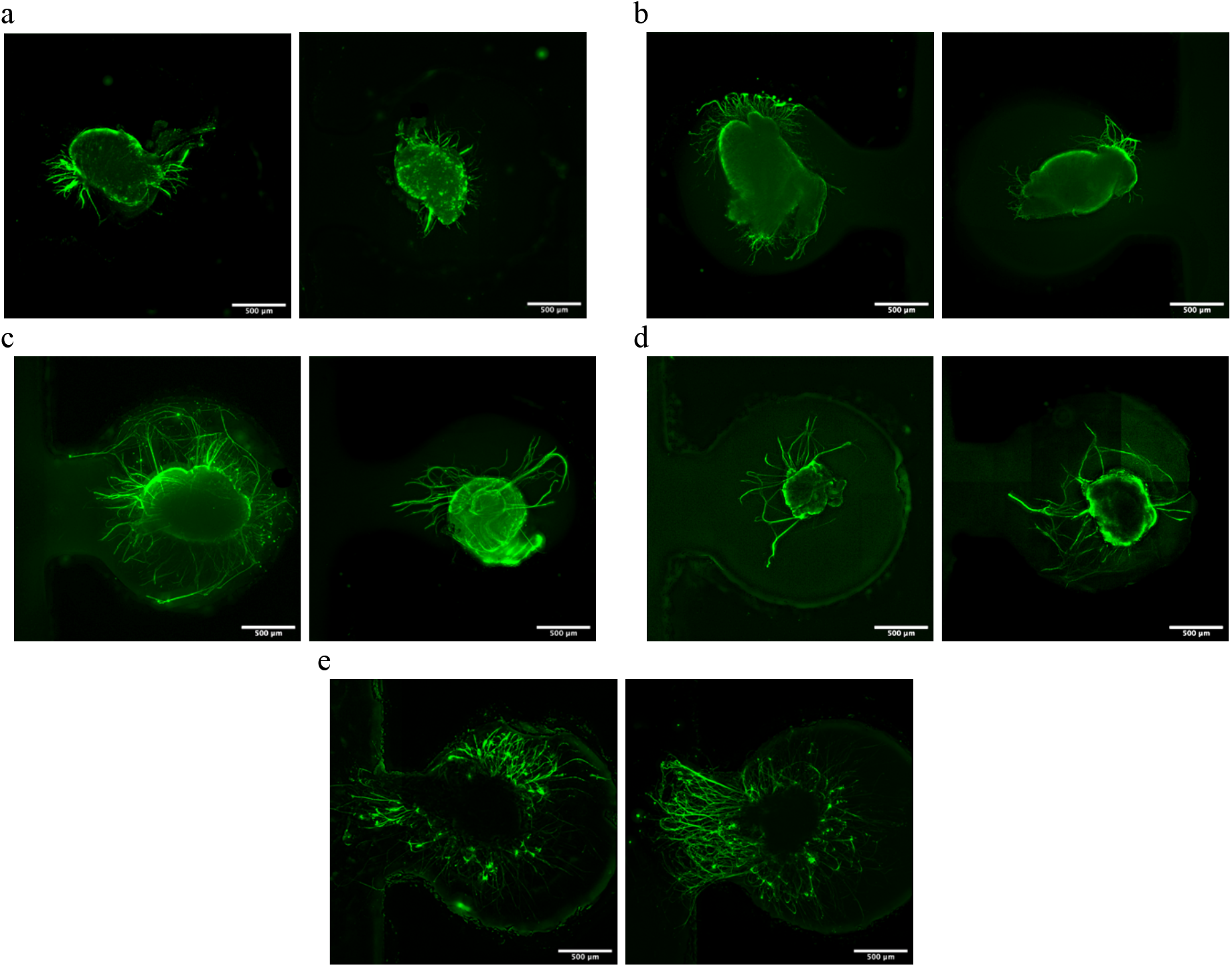
Representative images of DRGs in 3 % GelMA 95DoF in PBS (a), HBSS (b), BM-PHR (c), BM+10mM GSH (d), and Ultimatrix (e). Scale bar is 500μm. Brightness and contrast of the images were enhanced, and background noise was removed through ImageJ software for demonstration purposes.

### 3.4. Relationship between GelMA hydrogel stiffness and DRG neurite outgrowth

Linear regression has revealed a strong relationship between the GelMA hydrogel G’ value and mean neurite length with increased storage modulus resulting in shorter neurite lengths (Figure 13). Hydrogel storage modulus was a significant predictor of mean neurite length, explaining 83.1 % of the observed variation in mean neurite length (R^2^ = 0.831, F(1,4) = 19.63, p<0.0114).

**Figure 13.**
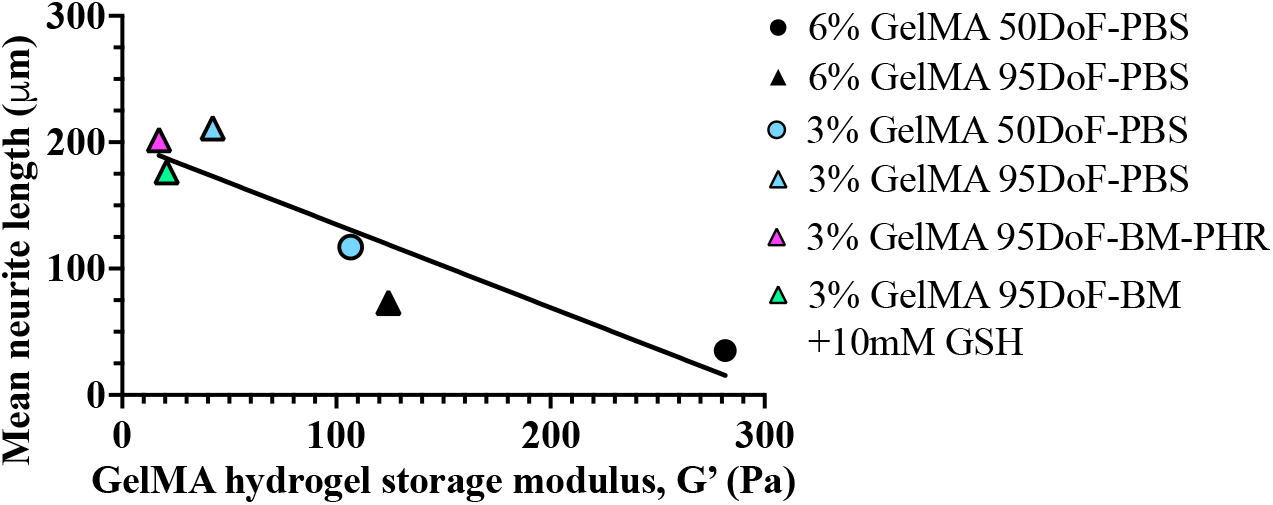
Simple linear regression of mean neurite length as a function of storage modulus values across tested hydrogel formulations.

## 4. Discussion

### 4.1. Hydrogel physical properties

#### 4.1.1. Effect of GelMA concentration

Higher concentrations of GelMA offer more methacryloyl (MA) groups for polymerization, resulting in the formation of dense polymer networks reflected in higher stiffness values and reduced swelling of hydrogels [6,35]. The availability of functional groups also impacts the dynamics of the polymerization reaction [35]. We have observed that for hydrogels prepared in BM, decreasing GelMA concentration from 6 % to 3 % leads to an increase in SF for both 50 DoF and 95 DoF GelMA, indicating slightly higher percentage of uncrosslinked polymer leftover after polymerization in the case of 3 % GelMA-BM compared to 6 % GelMA-BM (Figure 2). For hydrogels prepared in PBS or BM-PHR, negligible differences in SF between 3 % and 6 % GelMA were observed indicating that at the given light intensity and exposure time, maximum polymerization has been achieved for both concentrations tested. These findings also point out that application of BM leads to insufficient crosslinking compared to PBS or BM-PHR, which will be discussed in detail below.

In general, average SF values for 6 % and 3 % GelMA hydrogels were lower compared to those reported by Lim et al. for 10 % GelMA hydrogels prepared using the same solvent (PBS) and concentrations of Ru/SPS (0.2/2 mM), which could be explained by the higher visible light intensity used in our study (63.7 mW/cm^2^) compared to 30 mW/cm^2^ used by Lim et al. [8].

Further, a significant increase in the SReq for 3 % GelMA-PBS hydrogels compared to 6 % GelMA-PBS 50 DoF and 95 DoF indicates a formation of loose polymer network able to retain more liquid (Figure 3 (a)). Further rheological examination revealed a decrease in stiffness for 3 % GelMA-PBS hydrogels compared to 6 % GelMA-PBS hydrogels for both DoF, confirming lower crosslinking density reflected in higher SReq values (Figure 5 (a)). Overall, both GelMA concentrations explored demonstrated storage moduli below 1 kPa while maintaining efficient crosslinking (SF < 30 % as indicated by Lim et al [8].

#### 4.1.2. Effect of GelMA DoF

The degree of functionalization (DoF) of GelMA (%) is defined as the percentage of gelatin amino groups, primarily those on lysine residues, that are functionalized with methacryloyl (MA) groups, thereby determining the density of photocrosslinkable sites within the polymer [5,36]. Consequently, increasing the DoF increases the number of photocrosslinkable functional groups available during gelation, which can promote the formation of a more densely crosslinked polymer network under equivalent photocrosslinking conditions. For GelMA hydrogels prepared using UV-light-sensitive photoinitiators such as Irgacure 2959, high DoF (> 95 %) GelMA produces stiffer hydrogels compared to medium DoF (50 %) GelMA [4]. In the work by Walsh et al., no significant difference in storage moduli has been reported between the GelMA of medium and high DoF [17].

In this study, we have explored the differences in physical properties between medium (50) and high (95) DoF GelMA hydrogels prepared while maintaining a constant Ru/SPS concentration and blue light illumination settings (intensity and time). Unexpectedly, we observed that 60s of blue light exposure with unchanged intensity resulted in the formation of hydrogels with higher G’ values when 50 DoF GelMA was used compared to 95 DoF GelMA (Figure 5 (a)). This finding was consistent for both GelMA concentrations explored (3 % and 6 %). While high G’ values indicate increased stiffness and crosslinking density, this was not reflected in the swelling ratio values, with 3 % hydrogels demonstrating no significant difference in SReq between 50 DoF and 95 DoF, and 6 % GelMA 95 DoF hydrogels demonstrating lower SReq values compared to 6 % GelMA 50DoF (Figure 3 (a)).

Notably, 6% GelMA hydrogels of either DoF demonstrated no significant increase in the storage modulus when the light exposure time was increased from 60 to 90 and 120 s, indicating that sufficient crosslinking was achieved within just 60 s (Figure 6). Additionally, no statistically significant difference in the SF values between 50DoF and 95DoF GelMA was observed for both 3 % and 6 % polymer concentrations (Figure 2). The lower swelling ratio of 6 % GelMA 95DoF hydrogels could indicate heterogeneous crosslinking density rather than insufficient crosslinking.

#### 4.1.3. Effect of the solvent

Because the primary objective of this study was to develop GelMA formulations that support 3D neuronal growth in vitro, we investigated whether the medium used for hydrogel preparation and photocrosslinking influenced cell viability. PBS is most commonly used for GelMA hydrogel fabrication [4,13,14], but DMEM has also been utilized to create 3D scaffolds for cell culture [37,38].

We have observed that for low concentration of GelMA (3 %) reconstituted with BM (without phenol red) results in the incomplete crosslinking indicated by high SF values. Some of these samples have completely dissolved in PBS at 37 °C overnight, which is shown as 100 % SF (Figure 2). Although no significant difference was observed in SF values of 6 % GelMA hydrogels between the PBS and BM samples, there was a noticeable increase in SReq for BM samples compared to PBS, indicating the formation of less dense polymer network (Figure 3 (b)).

These findings are in agreement with previous reports on the contribution of media components to photopolymerization reaction kinetics. For example, in the study by Pamplona et al., GelMA hydrogels prepared with PBS have demonstrated significantly higher stiffness measured by AFM and higher MA conversion rate compared to the samples prepared in DMEM [37]. According to the study by Monfared et al. on photopolymerization kinetics of *N,N*-dimethylacrylamide (DMA) in cell culture media using UV-light, individual DMEM components were shown to inhibit polymerization [38]. Amino acids and vitamins such as L-cysteine and pyridoxine interfere with photopolymerization through two mechanisms: (i) inhibition of polymerization initiation through radical scavenging, and (ii) premature termination resulting from transfer of the propagating radical chain end onto media species [38].

Interestingly, the radical scavenging effect of media components was not as prominent when the BM with phenol red was utilized for polymerization compared to BM without phenol red. 3 % GelMA-BM-PHR hydrogels have demonstrated statistically significantly lower SF values compared to BM, but still higher compared to PBS.

Therefore, we hypothesized that phenol red in DMEM plays a role in facilitating photopolymerization of GelMA in DMEM. There are limited reports in literature on the application on phenol red for photopolymerization. The other studies on the impact of DMEM on photopolymerization which were referenced above have utilized phenol red-free DMEM. In the study by Lee et al., GelMA was mixed in with the phenol-red containing DMEM and 3D-printed, and the distribution of the pink colour intensity on the cross-section of the printed structure served as an indicator of higher and lower crosslinking density [39]. The authors have not provided the motivation behind using phenol red colour intensity as crosslinking efficiency indicator but have stated that the colour has faded when the higher UV-light exposure doses have been used. This could indicate that facilitated crosslinking rates have led to structural changes in the phenol red molecule.

Another study by Sundarakrishnan et al. has reported that phenol red has accelerated polymerization of silk tyrosine and resulted in the formation of stiffer material through the formation of the covalent bond between phenolic hydroxyl group and tyrosine [40]. It is possible that in the case of GelMA photopolymerization, phenol red undergoes oxidation and either (i) reacts with media species preventing them from attacking the growing radical chain; (ii) forms additional covalent bonds between the MA and phenolic hydroxyl groups. Although the addition of phenol red improved the crosslinking efficiency of GelMA hydrogels prepared in DMEM, it also has several drawbacks, including weak estrogenic activity and quenching of fluorescence from fluorophores emitting in the 400-500 nm range. These limitations should be considered when designing GelMA hydrogels for 3D cell culture and in vivo applications [40].

#### 4.1.4. Effect of GSH addition

Antioxidants, specifically GSH, have been previously used with GelMA to improve hydrogel biocompatibility and survival of encapsulated cells. For example, in the study by Lin et al., treatment of cells with GSH prior to their encapsulation in GelMA has resulted in increased cell proliferation rates and viability [41]. In the study by Wang et al. GelMA hydrogels manufactured in the presence of GSH have shown good biocompatibility and improved cell resistance to oxidative stress, however, physical properties of hydrogels remained unchanged when GSH was added or chemically grafted on to the GelMA polymer [42].

As discussed above, 3 % GelMA hydrogels prepared with BM have demonstrated poor crosslinking and high SF values. We have observed that upon the addition of GSH, SF values of 3 % GelMA-BM hydrogels have significantly decreased, indicating improved crosslinking (Figure 4). This finding was consistent for both 50DoF and 95DoF GelMA hydrogels. Additionally, while we were not able to examine the rheological profile of 3 % GelMA 95DoF-BM hydrogels, 3 % GelMA 95DoF-BM hydrogels prepared with 5 and 10mM of GSH demonstrated storage moduli values close to those of 3 % GelMA 95DoF-PBS hydrogels (Figure 5 (b)).

Interestingly, upon the addition of GSH to PBS or BM-PHR for crosslinking, no difference in SF was observed compared to samples crosslinked without GSH (Figure 4). As discussed above, free radical scavengers in BM interfere with different steps of photopolymerization process. In the case of Ru/SPS-mediated crosslinking we can identify the following steps: (i) Ru^2+^ is being photoexcited and donates electrons to SPS, (ii) SPS dissociates into sulfate radicals, (iii) radicals in turn initiate radical chain-growth polymerization of GelMA (31,39). BM species can interfere with any of these steps by: (i) accepting electrons from Ru and preventing sulfate radical formation, (ii) reacting with generated sulfate radicals, (iii) reacting with growing radical chain. GSH has an active -SH functional group performing as an electron donor. We hypothesize that GSH can prevent the unwanted side reactions by neutralizing oxidizing media species, but also by participating directly in the formation of the radical chain. The latter event can occur through the radical chain transfer, where during photopolymerization a propagating radical chain extracts hydrogen from GSH transforming it into the thiyl radical which reacts with MA double bond. This leads to the formation of a new radical containing covalent bond with GSH which can continue to propagate, or reaction termination occurs.

### 4.2. Neurite outgrowth from DRG explants in GelMA hydrogels

As discussed above, GelMA hydrogels are widely used for cell encapsulation and have been shown to support the growth and proliferation of various cell types due to the abundance of RGD cell-adhesive motifs [10]. However, different cell types have different needs when it comes to the biomaterial design in terms of substrate stiffness and bioactivity. Here we discuss GelMA hydrogel design considerations and modification strategies which can be used to improve the growth of neurons. The zero-growth fraction and neurite length were considered complementary measures of hydrogel performance. Previous studies have reported that a proportion of harvested DRG explants may fail to exhibit neurite outgrowth irrespective of culture condition, reflecting variability introduced during tissue isolation and preparation [43]. In the present study, however, the proportion of zero-growth explants also varied among GelMA formulations, suggesting that hydrogel properties may contribute to whether detectable neurite outgrowth is established. Such formulation-dependent effects could arise from differences in the physical properties of the hydrogel network, cell-matrix interactions, or cellular responses to the photocrosslinking environment. Reporting the zero-growth fraction separately therefore captures an aspect of hydrogel performance that would be lost if analysis was restricted to neurite length among explants that successfully initiated outgrowth. Among these growing explants, neurite length subsequently provides a complementary measure of the extent to which the hydrogel supports neurite extension.

In our study, we have observed that the hydrogels that have supported the most incidence of neurite outgrowth from DRGs (the lowest % of zero-growth) contained 3% GelMA 95DoF polymer (Table 3). Further, from statistical analysis it was evident that decreasing GelMA concentration from 6 % to 3 % and increasing GelMA DoF from 50 % to 95 % were effective strategies for increasing the median length of axons (Figure 9). These modifications were also shown to decrease the storage modulus of the hydrogel (Figure 13 (a)). Linear regression model has revealed a strong correlation between the stiffness of the hydrogel and neurite length (Figure 13). Notably, all hydrogels used for DRG encapsulation have demonstrated storage modulus values <1 kPa. However, 3 % GelMA 95DoF formulations (BM-PHR, PBS, HBSS and BM+10 mM GSH) which exhibited the highest median neurite length values all have presented with G’ below 100 Pa which is likely to be the upper G’ boundary allowing for 3D growth of DRG neurites.

It has been widely recognized that neurons prefer softer substrates to grow in with stiffness values close or matching to those of the brain and spinal cord tissues in the range of 0.1-3 kPa [44–46]. A review by Samanipour et al. has discussed that softer substrates (100-500 Pa) provide biophysical cues for differentiation of neural and mesenchymal stem cells into neurons, while stiff materials (1-10 kPa) drive predominantly glial lineage differentiation [47]. Similarly, a recent study reports that neural progenitor cells demonstrate higher rates of neurite outgrowth in softer hydrogels with G’ = 200 Pa [48]. Another study suggests that while CNS neurons demonstrate greater extent of neurite sprouting and branching on softer substrates, they can also extend axons on stiffer gels through cellular connections with astrocytes which prefer stiffer materials [49]. In the study by Ribeiro et al. DRG neurons demonstrate higher degree of mechanosensitivity compared to CNS neurons, preferring to grow on substrates with intermediate Young’s modulus of ∼1 kPa, however, only 2D outgrowth was examined [50]. The study by Man et al. has showed that 3D outgrowth of DRG neurites was found to be directly dependent on the stiffness of fibrin gels, with gels that have low compressive modulus (<3 kPa) demonstrating neurite lengths between 200-250 µm [32].

When cells or tissue explants are encapsulated directly within a photocrosslinkable polymer such as GelMA, they are exposed to multiple components of the photopolymerization process. The generation of free radicals during photocrosslinking, as well as reactive byproducts, can adversely affect cell viability [51,52]. Because most cell culture media contain vitamins and antioxidant molecules, preparing hydrogels in culture medium has been proposed as a strategy to improve the viability of encapsulated cells by reducing oxidative stress during photocrosslinking. However, our study supports the claim that cell culture media components directly hinder photopolymerization process, leading to insufficient crosslinking and loose hydrogel network formation. Application of BM in the absence of phenol red and GSH failed to form the hydrogel, and all explants were lost in the experiment for this group. On the other hand, formation of GelMA hydrogels in BM with phenol red was successful and all DRG explants have demonstrated high median neurite length values, although not significant from the PBS group of the same GelMA concentration and DoF (Figure 11). HBSS was tested as a substitute for PBS for 3 % GelMA 95DoF fabrication due to its high glucose content, which was hypothesized to improve DRG viability, however, no significant difference was found between the two or with BM-PHR group. Similarly, an addition of antioxidant GSH for 3 % GelMA photopolymerization in BM didn’t demonstrate a significant improvement in neurite outgrowth from DRGs compared to non-supplemented PBS formulation, although it played a significant role in achieving efficient crosslinking. Overall, while different solvents didn’t demonstrate a significant impact on DRG neurite outgrowth, the choice of a solvent for hydrogel fabrication should be carefully considered with regards to photopolymerization efficiency.

Overall, DRG explants exhibited longer neurite outgrowth in the Ultimatrix control hydrogel than in any of the GelMA formulations tested (Figure 11). Ultimatrix is a commercially available basement membrane extract widely used as a three-dimensional cell culture matrix and scaffold for organoid culture. Derived from the mouse Engelbret-Holm-Swarm (EHS) tumor, it contains numerous extracellular matrix (ECM) components, including laminin, collagen IV, entactin, and associated growth factors, which can provide biochemical cues that promote neurite extension. While Ultimatrix closely recapitulates aspects of the native extracellular environment, its tumor-derived origin, batch-to-batch variability, limited compositional control, and challenges in large-scale manufacturing reduce its suitability for reproducible in vitro studies and translational in vivo applications [53]. GelMA offers a highly tunable semi-synthetic matrix and can be further modified to achieve the biological complexity of cellular environment through the addition of ECM components and growth factors. Future studies will investigate the addition of such factors like laminin or laminin fragments [54], as well as glial cell derived neurotrophic factor (GDNF) [55].

## 5. Conclusions

In this study, we demonstrated that GelMA hydrogels photocrosslinked using visible blue light provide a suitable platform for the encapsulation and three-dimensional growth of primary DRG explants. The crosslinking efficiency, network density, and mechanical properties of the hydrogels could be systematically tuned by varying GelMA concentration and degree of functionalization (DoF). We showed how using different solvents to reconstitute GelMA affects photopolymerization efficiency, and how these effects can be averted by addition of phenol red or GSH. GelMA hydrogels have supported the outgrowth of axons from encapsulated DRG explants which was analyzed using the custom-developed semi-automated image analysis tool. Further, comprehensive statistical analysis established quantitative relationships between GelMA formulation, hydrogel mechanical properties, and DRG neurite outgrowth. Among the formulations examined, 3 % GelMA with a 95 % degree of functionalization and a storage modulus below 100 Pa supported neurite outgrowth, providing a rational basis for engineering photocrosslinkable hydrogels tailored to neural tissue models and regenerative medicine applications.

## Supporting information

Supplemental Section

## 6. Acknowledgements

The authors are grateful to work and live on the traditional, ancestral and unceded territory of the Coast Salish Peoples, including the Musqueam and Squamish First Nations. The authors would also like to acknowledge fellowship support through UBC Centre for Blood Research Graduate Award Program and UBC Elwyn Gregg Memorial Scholarship, The UBC Applied Statistics and Data Science (ASDA) group for statistical consultation, Super Resolution Microscopy Core at UBC for imaging support, and Dr. Wolfram Tetzlaff research group at the International Collaboration for Repair Discoveries (ICORD) at UBC. The authors would like to acknowledge the support of the Government of Canada’s New Frontiers in Research Fund (NFRF) [NFRFT-2020-00238], the Natural Sciences and Engineering Research Council of Canada (NSERC) and the Canada Foundation for Innovation (CFI).

