## Supplemental Section for "Strategies to improve neurite outgrowth from primary neurons in gelatin methacrylate hydrogels polymerized with visible light exposure inside a microscale 3D model"

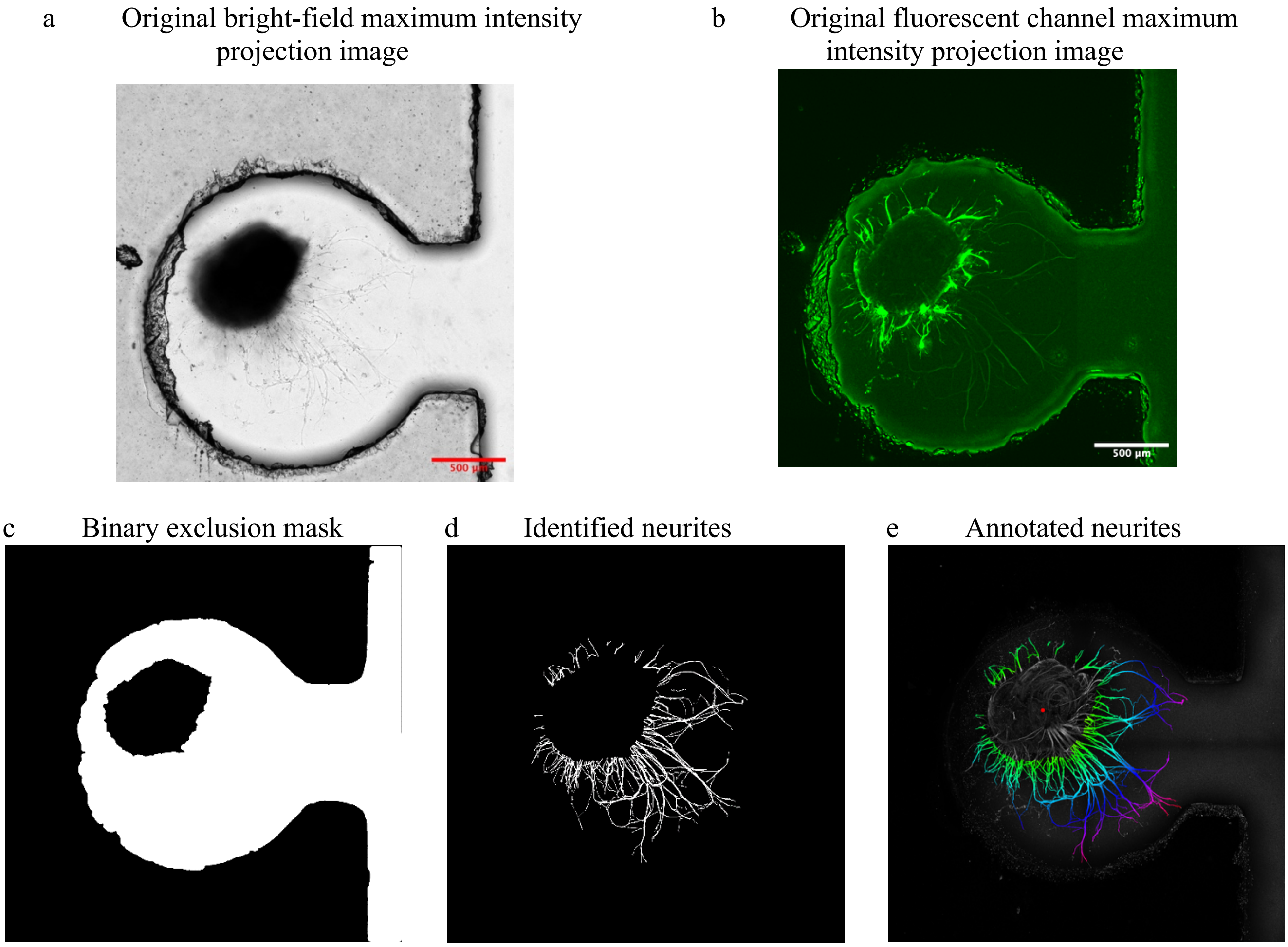


*Figure S 1. Examples of intermediate processing steps from a custom Automated Quantification Software for DRG neurite identification and length quantification. 1.a Shows a 2-dimensional Minimum Intensity Projection, computed from the Bright-Field Z-Stack, used to identify the boundaries of the DRG body and the interior region of the growth chip. 1.b Shows the 2-dimensional Maximum Intensity Projection of the fluorescent channel, corresponding to the location of the neurite extensions. The relatively high background intensity within the growth chip interior presents a challenge for segmentation. 1.c. Shows the inclusion mask defining the "valid region" in which neurite extensions are considered possible. This excludes the DRG body and the surrounding support material of the growth chip. 1.d Shows the resulting segmented neurite extensions, computed layer-by-layer from the fluorescent Z-stack before flattening. This pixel mask, along with the location of the centroid of the DRG body is used to compute a simplified Sholl analysis to provide an approximation of neurite length. 1.e shows a visualization of neurite length within the image, where the hue of a pixel varies with distance from the DRG centroid. This visualization is overlaid on a normalized, greyscale maximum intensity projection of the fluorescent stack to provide human-readable visualization of which image regions were identified as neurite for a posteriori interrogation of the reported neurite lengths.*
